# A climate-based model for tick life cycle: an infinite system of differential equation approach

**DOI:** 10.1101/2021.09.02.458669

**Authors:** Cyrine Chenaoui, Slimane Ben Miled, Mamadou Sadio Ndongo, Papa Ibrahima Ndiaye, Mourad Rekik, Mohamed Aziz Darghouth

## Abstract

The distribution of ticks is essentially determined by the presence of climatic conditions and ecological contexts suitable for their survival and development.

We have developed a general tick biology model to study the major trends due to climate change on tick population dynamics under different climate conditions.

We build a model that explicitly takes into account stage into each physiological state through a system of infinite differential equations where tick population density are structured on an infinite discrete set. We suppose that intrastage development process is temperature dependent (Arrhenius temperatures function) and that larvae hatching and adult mortality are temperature and precipitations dependent.

We analysed mathematically the model and have explicit the *R*_0_ of the tick population. Therefore, we performed a numerical analysis of the model under three different climate conditions (tropical, Mediterranean and subarctic climates) over the short term using climatic data from 1995 to 2005, as well as long-term simulations from 1902 to 2005.

## 1. Introduction

There are up-to-date more than 900 ticks species showing specific distributions over the world, and only 25 of these species are representing major threats to humans and animals, essentially through the transmission of pathogens[21]. Indeed, amongst all the hematophagus vectors, ticks are those transmitting the highest range of pathogenic, bacteria, parasites and viruses [35], causing important human and animal diseases such as for instance tick-borne encephalitis in humans and malignant theileriosis in livestock [3]. The distribution of ticks is essentially determined by the presence of climatic conditions and ecological contexts suitable for their survival and development, including the presence of the required hosts for the feeding of immature and adult stages. The distribution and population density of tick species as well as the relative importance of the transmitted tick-borne pathogens for humans and animals depend greatly on the suitability of the eco-climatic conditions regarding the tick biological requirements.

Tick biology is complex and exhibit a high level of biological polymorphism observed between the different tick genera and even between the different species within a given genus. For instance, in the *Hyalomma* genus, some species are relatively less adapted to drought and high temperature, like for instance *H. lusitanicum* and to a lesser extent *H. scupense*, comparatively to *H. dromedarii* and *H. anatolicum* [38, 8]. This high polymorphism is exacerbated by the complexity of the interaction between the ticks, their hosts, and the pathogens they transmit. Accordingly, understanding and predicting the mechanisms leading to a determined phenology is quite impossible. The ongoing dynamic of climate change is bringing an additional layer of variability that imposes to adopt a proactive attitude to those concerned by on the control of the major ticks and ticks borne diseases (TBD) in humans and animals. In this changing context, where working as usual is certainly not the solution, predicting the size of tick populations and anticipating the impact of different control actions remain difficult. However, modeling represents a powerful alternative that could overcome these difficulties, offering then valuable decision aid data that bring a rational basis for deploying proactive measures against ticks and TBD.

The object of the present work is to develop a general tick biology model for analysing the major effects of climate change on the evolution of tick population dynamic in different climate conditions (tropical, Mediterranean and subarctic climates).

Models for tick population dynamics often describe in a discrete way the various stages of ticks development from egg-larva-nymph-adult (*e.g*. [9]), whether the ticks are attached to hosts [20], and if disease is part of the model, whether the ticks themselves are infected [19, 10]. For instance, [31, 24, 22] and more recently Lou and Wu [17] proposed a model where ticks are subdivided in the three stages (larvae, nymphs and adults) with stage progression only through a blood meal on a vertebrate host (two types of which are considered in the model).

Empirical studies confirm that the developmental and questing activity of ticks are regulated by the climate [16, 27, 28, 32, 5, 26]. In addition, seasonal variations are one of the major factors in the transition from one stage to another [6]. Given the critical impact of climate change on tick life cycles, several models has been constructed [7, 31, 25, 2, 4].

[31] proposed a model that considers diapause and density-dependent regulation of nymphs and adult stages, as well as temperature-dependent rates of development and egg production, and density-independent but climate-dependent mortality rates for both larval and adult stages. Their model accurately predicts both the seasonality and the annual spectrum of variation in life stages density tick species *Rhipicephalus appendiculatus*. More recently, [11] developed an age-structured model to examine the effect of shifts in average temperature on seasonal activity of ticks and on interstadial development.

Also, Ogden et al. [25] developed a dynamic population model of *Ixodes scapularis*, where tick development rates are temperature-dependent time delays, calculated using mean monthly normal temperature data and then applied to forecast theoretically the physical boundaries for *Ixodes scapularis* establishment in Canada and host-finding success.

To predict the possible consequences of climate change on ticks’ dynamics, a more extensive understanding of the relative contributions of temperature-dependent mechanisms is crucial. As an extension of previous work [25], we propose in this study to model on a temperature-dependent basis the dynamics of tick populations using a system of finite non-autonomous differential equations where tick population density are structured on a finite discrete set. Temperature is the most extensively analysed climatic factor for establishing the effect of the phenomenon of climate change allowing therefore to extend its application for exploring the putative effects of changing climatic conditions on tick dynamics using the temperature-dependent model developed in the present work.

We performed a numerical analysis of the model under the Mediterranean climate of north-east Tunisia, focusing on the endemic tick species *H. scupense*. We also extended our analysis to two other climates, namely, a tropical climate in Senegal with the tick *Amblyomma variegatum*, and a subarctic climate in Canada with the tick *Ixodes cookei*. In order to evaluate the predictive functionality of the model we also run, in each of the three climatic contexts with the corresponding endemic tick species, a simulation using climatic data to assess the short-term trends during the decade 1995 − 2005 and the long term trends over one century from 1902 until 2005.

In section 2, we describe the mathematical model. Later in section 3 we exhibit numerical result respectively in constant and fluctuating temperature. Finally, in 4 an analysis of results is presented, and the limits of our model are discussed.

## 2. Model description

Under the Mediterranean climate of North Africa [8], *H.scupense* adults take a blood meal during the summer. Females detach from the animal, layout eggs and die. The larvae start questing for a host for feeding and then moult into nymphs. The nymphs, also, take one blood meal and detach. Immature ticks are observed on animals during the autumn. The engorged nymphs look for a shelter (wall cracks and crevices in the barn) and enter diapause (see figure 1).From the end of the spring to the beginning of summer, the nymphs moult into adults and leave the shelters and start questing for a host.

**Figure 1:**
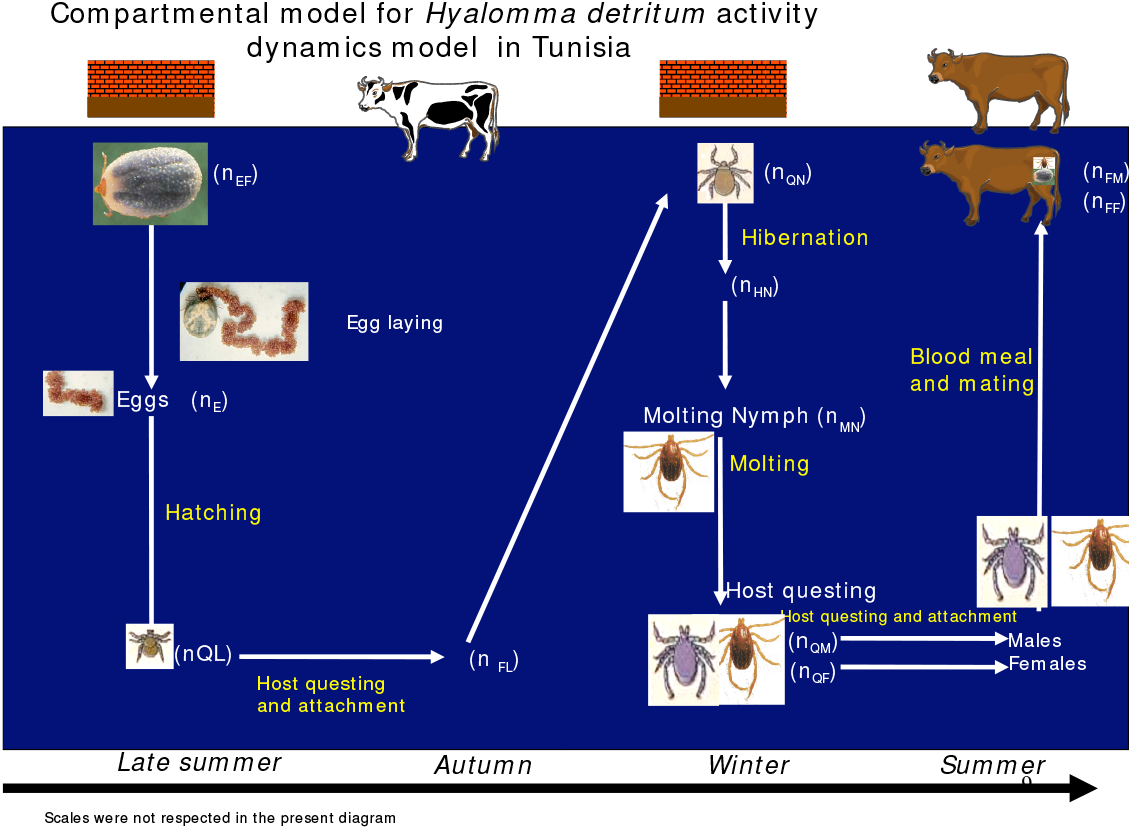
Different states of the tick

Thus, the physiological growth inside in stage is a continuous process, which can be approached by a physiologically structured model using Usher’s matrix **??**. The equation considered in this model is derived from a discrete physiological-structured model for a population in which individuals change their stage according to seasonality.

To illustrate the ideas underlying the model, we consider a population divided into four physiological stages, eggs, larvae, nymphs and adults, each one being structured by development stage, *s*. We suppose that *s* ∈ {*s*_*j*_ */ j* ≤ *p* − 1}, with *p* the stage number and denote for all *i* ≤ *p, e*_*i*_(*t*) the density of egg, *l*_*i*_(*t*) the density of larvae, *n*_*i*_(*t*) the density of nymph and *a*_*i*_(*t*) the density of adult at time *t* and in the development state *s*_*i*_. Capitals, *E*(*t*), *L*(*t*), *N* (*t*) and *A*(*t*), denoted the total population number at time *t*.

Let’s note, for every physiological stage *s*_*i*_ with *i* ≤ *p*, by 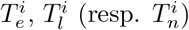 the interstate transition rate to the larva, nymph, and adult stages. We denote, 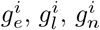 and 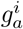 the intrastate transition rate corresponding to the development stage denoted for the egg, larva, nymph, and adult. It represents the physiological maturity increasing rate.

We also represent, *β*^*i*^, the eggs’ production rate and 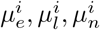 and 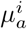thedeath rate of the egg, larva, nymph and adult at stage *s*_*i*_.

To simplify the model and as our objective is to study the effect of temperature on the life cycle, we will neglect the effects of density dependencies especially on nymph and larva mortality, as well as on transition rates. It is noted that the effect of density dependence has been discussed at length by different authors [12, 31, 20, 34, 9, 33, 23].

According to the above arguments, the model equations are (see figure 2):

**Figure 2:**
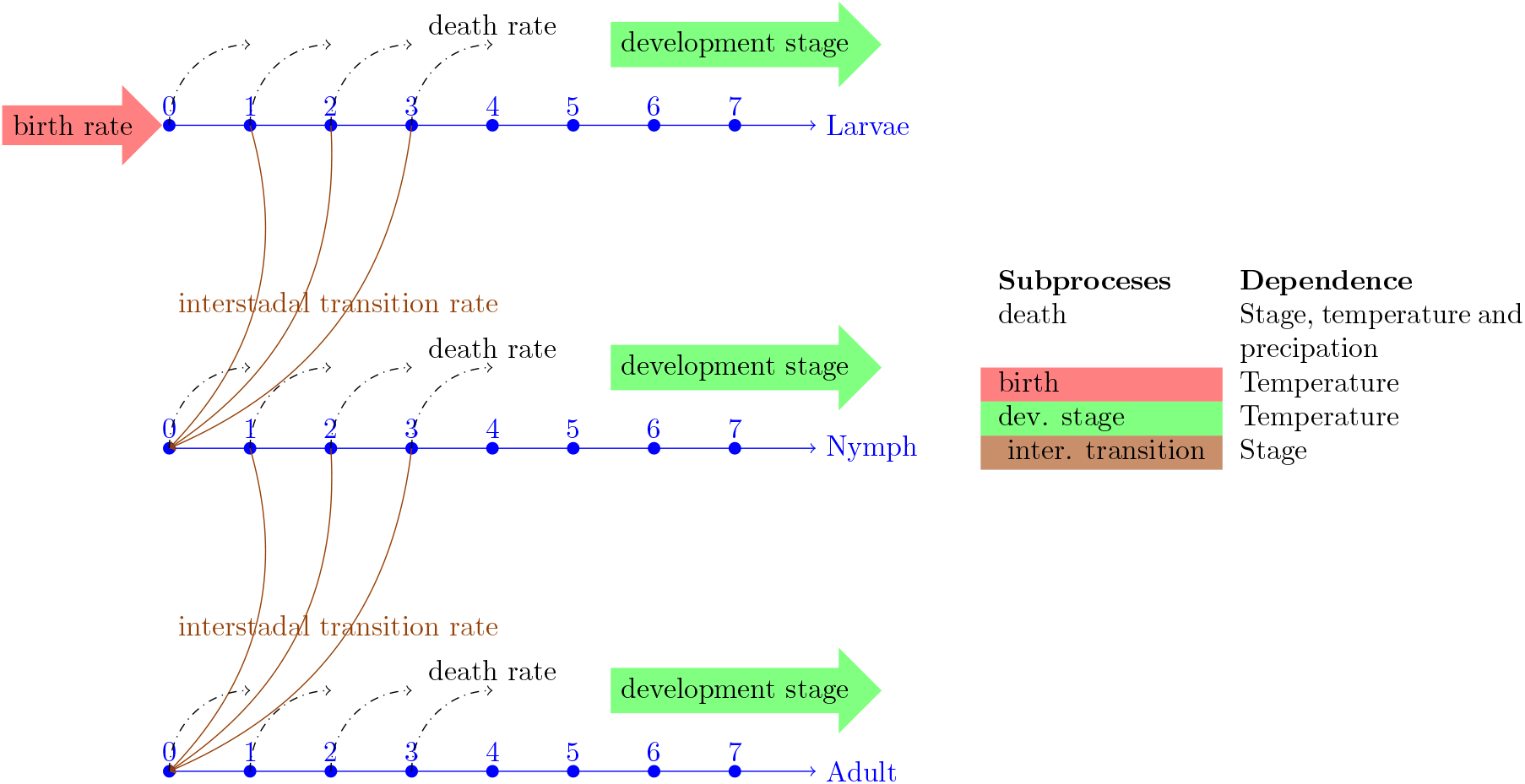
A schematic diagram for the model, identifying the input parameters and the model output (predicted number of ticks at each stage)

**Egg:**

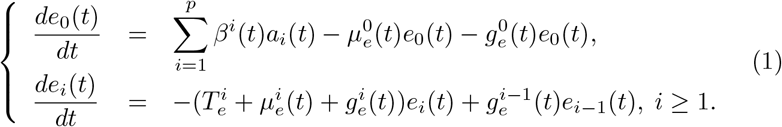

**Larva:**

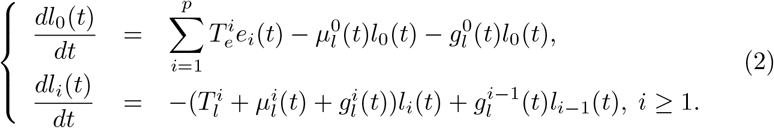

**Nymph:**

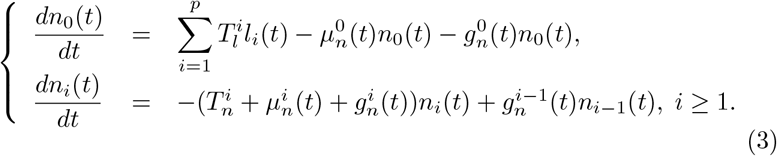

**Adult:**

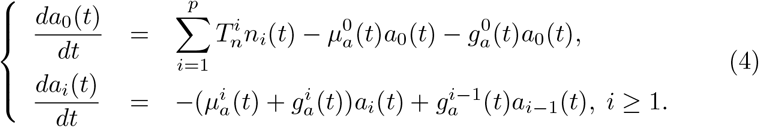

### 2.1. Temperature effect

In endothermic species (*e.g*. ticks), metabolic rates are governed by biochemical reactions and thus are temperature-dependent. Based on [15], growth, in particular, corresponds to the conversion of absorbed energy to structure and so is temperature-dependent along with other factors, then we could consider the metabolic rate as temperature-dependent. Since they are controlled by enzymes that are inactive at extreme temperatures, metabolic rates are reduced at low and high temperatures.

For simplicity, we suppose that only transitions rates, 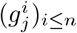, with *j* ∈ {*e, l, n, a*} are temperature-dependent, *i.e*. 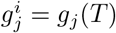 for all *i* ≤ *n*.However, as we suppose there is density dependence (*e.g*. no competition for blood meal), we will consider that interstate transition rates, 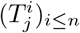, with *j* ∈ {*e, l, n, a*} depend only on physiological stage, (*s*_*i*_)_*i*≤*n*_, as following:

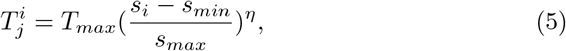

with *j* ∈ {*e, l, n*}, *η* ∈ ℝ_+_ a physiological parameter and *T*_*max*_ the maximum transitions rate. We suppose that *s*_*min*_ and *s*_*max*_, the min and max development stage for each physiological stage values are set at 0 and 1.

We have considered in our model that larva and nymph mortalities are constant in the tolerance temperature range, 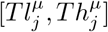 for *j* = {*e, l, n, a*} Randolph [29]. Therefore, let:

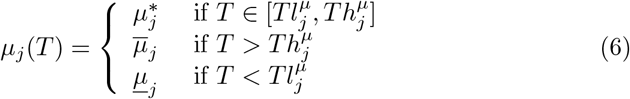

and

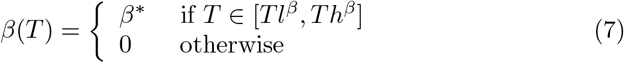

With, 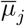 and <u>*μ*</u>_*j*_ the mean death rate at high and low temperature.

However, we suppose that egg hatching, larva, and nymph transition rates depend linearly on temperature *T* :

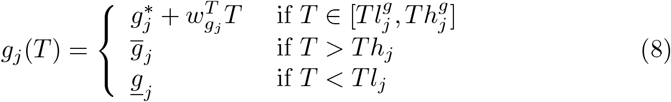

with for all 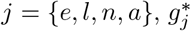 and 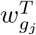, stand for the temperature effect of the intrastate transition rate (*i.e*. development rate) in the tolerance temperature range 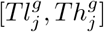. Overline and underline stand, respectively, for mean survival at high and low temperature. See table 1 for the definition of the parameter model.

**Table 1:** Input parameter values for the population model for the three sites: Canada (CA), Tunisia (TN) and Senegal (SN)

| | | $e$ | $l$ | $n$ | $a$ | Ref. |
| --- | --- | --- | --- | --- | --- | --- |
| Diapause temperature in Celsius, $Tl_j^g$, for the three sites: Canada, Tunisia and Senegal | | | | | | |
| CA |  | 14 | 12 | 4 | 5 | Unpublished data |
| TN |  | 14 | 12 | 4 | 5 | Unpublished data |
| CA |  | 14 | 12 | 4 | 5 | Unpublished data |
| Commun input parameter values for the three sites: Canada, Tunisia and Senegal |  |  |  |  |  |  |
| $g_j^*$ | Mean transition rate | -55 | -87.7692 | -6.87 | -20.07 | Unpublished data |
| $w_{g_j}^T$ | Effect of temperature, $T$, on transition rate | 6.06 | 6.8846 | 6.8846 | 6.8846 | Unpublished data |
| $\mu_j^*$ | Mean death rate | 0.0450 | 0.0159 | 0.0114 | 0.0196 | Unpublished data |
| $\bar{\mu}_j$ | Mean survival at high temperature in days | 3 | 3 | 3 | 3 | Unpublished data |
| $\bar{\mu}_j$ | Mean survival at low temperature in days | 25 | 25 | 25 | 25 | Unpublished data |
| $\beta_j^*$ | Mean egg production rate | | | | 2.1 | Unpublished data |
| $Th_j^\mu$ | Higher temperature tolerance in Celsius | 32 | 32 | 32 | 32 | Unpublished data |
| $Tl_j^\mu$ | Lower temperature tolerance in Celsius | -7 | -7 | -15 | -15 | Unpublished data |
| $Th_j^g$ | Higher temperature tolerance in Celsius | 30 | 30 | 30 | 30 | Unpublished data |
| $Th^\beta$ | Higher temperature tolerance in Celsius | 30 | 30 | 30 | 30 | Unpublished data |
| $Tl^\beta$ | Lower temperature tolerance in Celsius | 30 | 30 | 30 | 30 | Unpublished data |
| $\hat{g}_l$ | Maximum larva intrastadial transition rate | 100 | 100 | 100 | 100 | Unpublished data |
| $\hat{g}_l$ | Maximum larva intrastadial transition rate | 100 | 100 | 100 | 100 | Unpublished data |

## 3. Numerical results

We simulate ticks population of eggs, (*e*_*i*_(*t*))_*i*≤*p*_, larvae, (*l*_*i*_(*t*))_*i*≤*p*_, nymphs (*n*_*i*_(*t*))_*i*≤*p*_ and adults (*a*_*i*_(*t*))_*i*≤*p*_, densities for a finite physiological structure (*s*_*i*_)_*i*≤*p*_ (*p* = 100), and for three different climate conditions, tropical (Senegal), Mediterranean (Tunisia) and subarctic climates (Canada). We also performed simulations for two distinct periods: a 10-year period and a longer one of more than 100 years. Climate data used in this model are the Climatic Research Unit (CRU) TS Time Series datasets 4.03 [37]. The parameter values are set in tables 1.

### 3.1. Constant temperature simulation

We simulate the density of larvae, nymphs and adults for *T* = 24°C over a 25 month period. We note that the tick population at this temperature performs a little more than 2 cycles per year (see figures 3 and 4). Figure 4 indicates that the larvae are present at varying densities throughout the year at a constant temperature, with two peaks, one in March and the other in September and October. It’s important to note that the density of larvae is lowest in the summer (June and July) and winter (December and January). There is a slight overlap between the two stages for nymphs and adults (figure3), with two peaks after the preceding stage’s peak.

**Figure 3:**
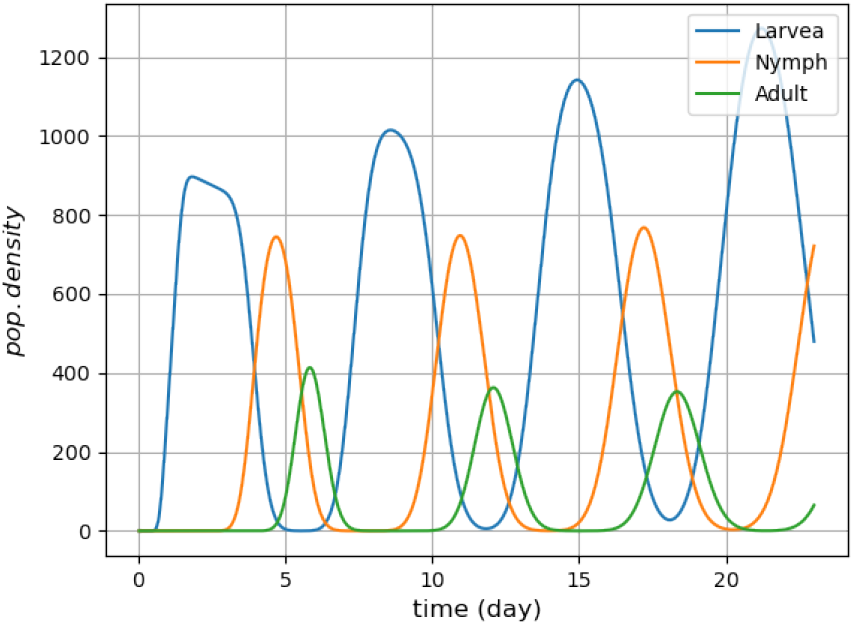
Larva, nymph and adult population densities

**Figure 4:**
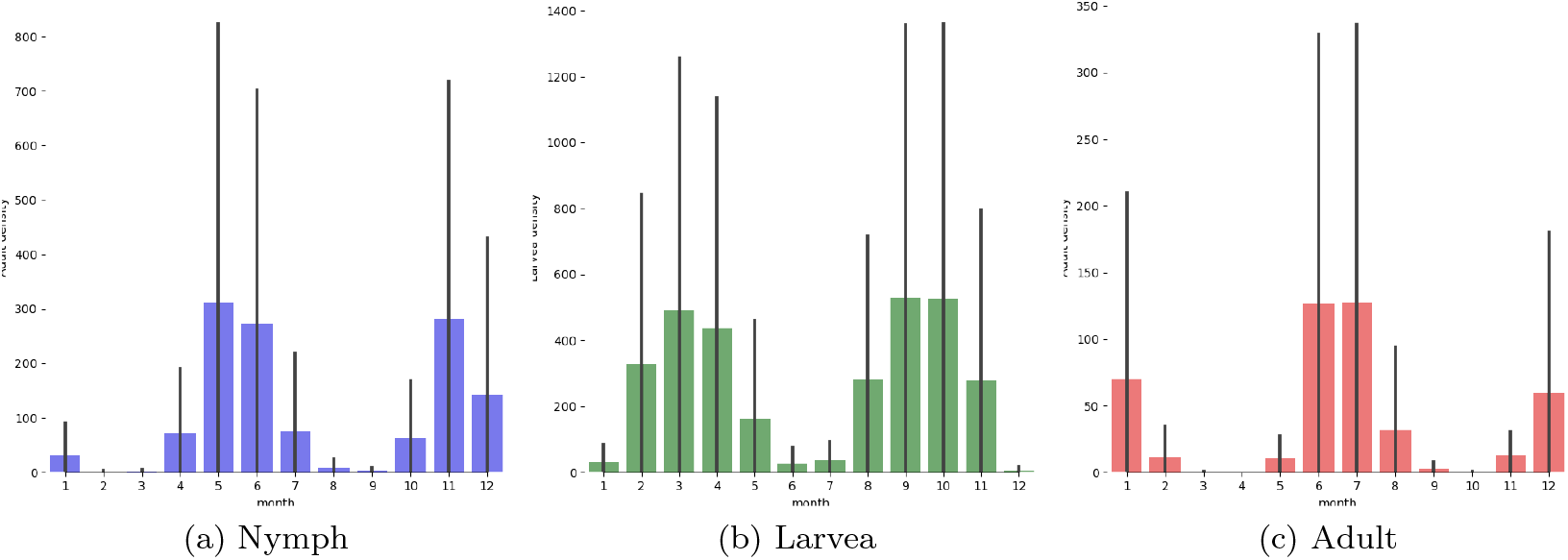
Larva, nymph and adult population densities by size

We then calculated the basic reproduction number *R*_0_ of the population for *T* ∈ ss[15, 33]°C. We notice that *R*_0_ admits a maximum at 22°C with a value of 3.4, then decreases slowly to *T* = 30°C (see figure 5a). The sensitivity analysis of the *R*_0_ (see figure 5b) in relation to the variation with respect to temperature *T* showed two sensitivity picks, one at 19°C and the second at 30.5°C. These two peaks correspond to the discontinuities of *R*_0_. Moreover, in the set [19, 30]°C, *R*_0_ is almost not sensitive to *T*.

**Figure 5:**
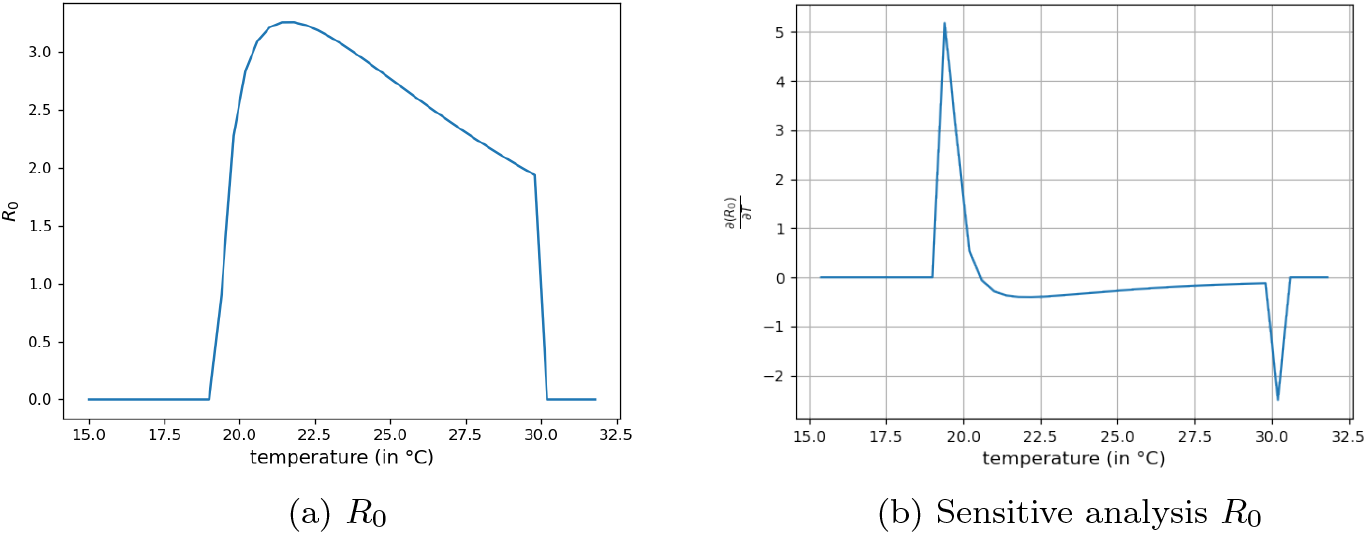
The basic reproduction number *R*_0_ by temperature

### 3.2. Simulations with fluctuating temperature

We performed a numerical analysis of the model under three different climate conditions (tropical, Mediterranean, and subarctic climates) and using climatic data from 1904 to 1995 (see figure 6). The physiological parameters of ticks have been set to the same values for three different climatic environments, except for diapause temperature that has been set about the country’s annual temperature distribution.

**Figure 6:**
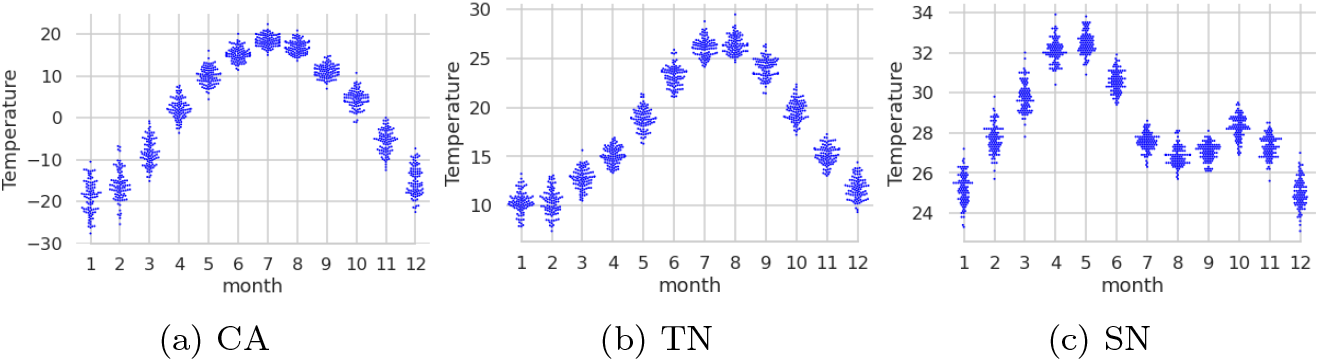
Monthly temperature distribution in the period 1904 *−* 1995 for Canada (CA), Tunisia (TN) and Senegal (SN)

Canada is characterized by cold and dry winters and warm (*i.e. T* = 20°C) summers (see figures 6a and 7a). The temperature between April to October is positive and does not exceed 20°C. The temperature decreases to − s20°C to − 30°C in January (see figure 6a).

**Figure 7:**
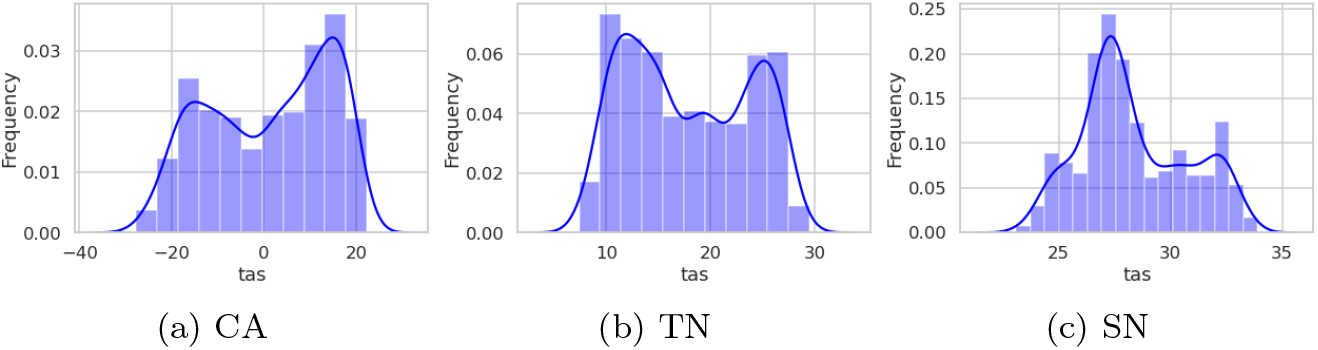
Temperature distribution in the period 1904 *−* 1995 and the period 1995 *−* 2005,respectively for Canada (CA), Tunisia (TN) and Senegal (SN)

Tunisia is characterized by a Mediterranean climate, mild (*i.e*. 10 °C) and wet winters and hot (*i.e*. 30 °C) and dry summers (see figures 6a and 7a).

Senegal is characterized by a dry period and a wet period (see figures 6a and 7a). The wet periods (July and August) are warm (Temperature 28 ° C) and the dry periods (March and April) are hot (see figure 6a).

#### Short-term simulations

First, we simulated the evolution of life cycles over 3 years in Tunisia and Senegal from June 2002 to June 2005. Given the extreme temperatures, the life cycle in Canada extends over 2 years, so we simulated the life cycle over 10 years from June 1995 to June 2005.

Our model predicts a significant difference in nymph and adult population density variations between the three countries (see figure 8). The amplitude of seasonal variations is observed in the Mediterranean climate, where the population density in Tunisia rises from 0 in winter to 2000 in summer (see figure 8b). As we move further south or north, the amplitude decreases; In Canada, we observe that seasonal fluctuations in adult density between 0 in winter and 800 in summer (see figures 8a).

**Figure 8:**
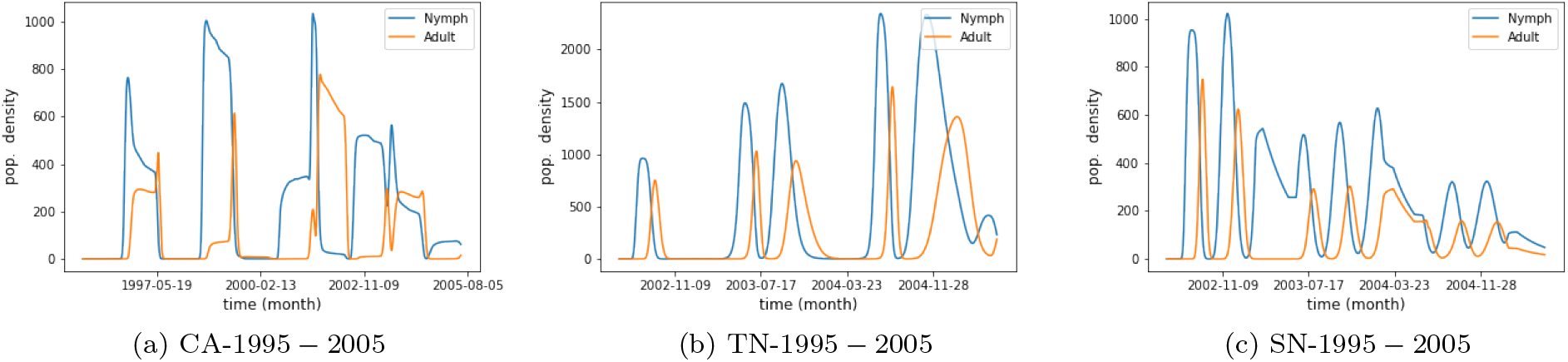
Total population density by countries and for the periods 1991 *−* 2015

In Senegal (see figures 8c) the population increases from 0 in winter to 1000 in summer.

We do notice, however, that the life cycle in Canada is around 2 years. In Tunisia, this duration is reduced to 1 to 2 yearly cycles, and in Senegal, it is reduced to 2 yearly cycles, with a significant overlap of nymphs and adults in some cycles.

#### Long-term simulations

We observe for the long term simulations, 1902 − 2005, that tick populations remain high over time in the three countries, indicating that a susceptible adaptation to only low temperatures is sufficient to keep ticks alive in a variety of different environments (see figure 9).

**Figure 9:**
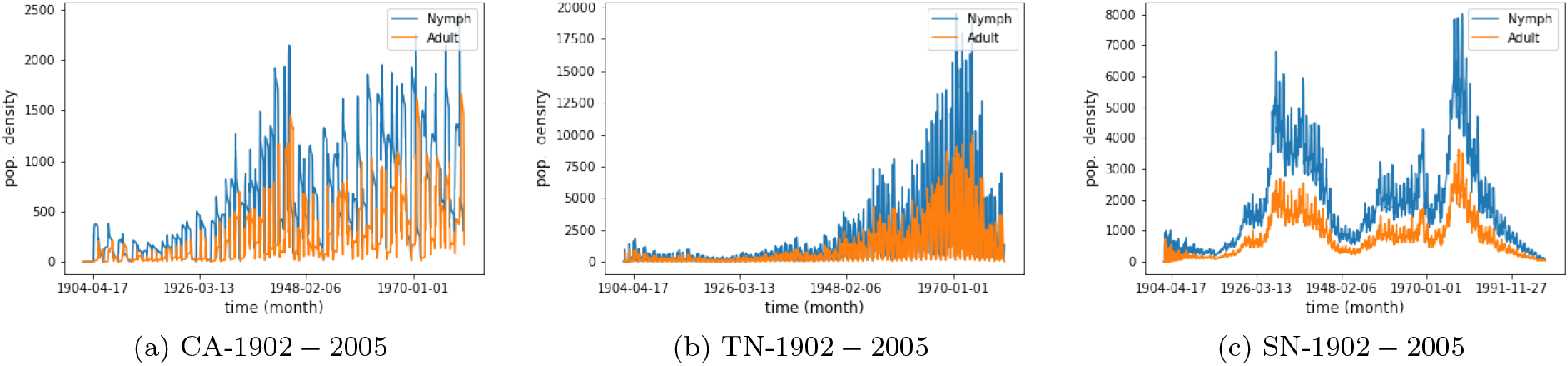
Total population density by countries and for the periods 1902 *−* 2005

Indeed, in Canada, the population density soars in the period 1940 − 45, then decreases sharply in late forties, after which it rises gradually before reaching a second peak in the latest ten years (figure 9a). In Tunisia, we also observe two peaks, a small one around the year 1960’s and a larger one in the 1970’s (figure 9b).

In Senegal, the first peak was observed in the period 1930 − 1940 (slightly earlier than in Canada) and the second around the 1980’s. There is also another smaller peak in the 1960’s (figure 9c). On the other hand, the fluctuations in density are smaller in Canada than in the other two countries.

## 4. Discussion and conclusion

We have constructed a model to study the dynamics of tick populations under the influence of a major parameter, namely temperature. This model is primarily intended to be used for assessing the overall effects of climate change scenarios on tick population dynamics. The dynamics of tick populations have been structured in three stages, assuming that each stage is subdivided in an infinite number of states composing an infinite discrete set. Therefore, tick population density satisfies a large set of differential equations. Our model is located at an intermediate level between the existing models assuming that physiological structuring is represented by a finite number of classes often small [30, 7, 14] and the continuous PDE models [18, 36].

Physiological models were developed by [18] where authors analyzed examples of a partial differential equation with physiological states belonged to an infinite non-countable set. However, the development of corresponding model equations is complex and, in many cases, does not allow displaying demographic parameters such as *R*_0_. Our approach succeeded in analytically exhibiting it.

Seasonality variation is particularly important to an organism with an annual life cycle. Analyzing the demography of these species is complicated as two time scales are involved, firstly within the year when the life cycle occurs and secondly, between years when reproduction and death occur. In several cases, the duration of the life cycle periods is directly related to temperature. The development rate evaluation methods require the calculation of the development fraction achieved on each day, a value depending on the daily temperature [7]. This calculation introduces complexity in the mathematical analysis. The originality of our model lies in considering temperature-dependent intrastate transition rate. This coupling approach enables us to solve the scale coupling problem without using two-time scales and/or delay to account for tick diapause, and therefore, highlight fluctuations in population density in correlation with seasonality.

We showed that seasonality is the main factor for the synchronization of different physiological classes (nymphs and adults) in the population. Indeed, we notice that in three climate conditions climates, both physiological stages are mainly asynchronous with a slight overlapping indicating the ongoing moulting process. We showed that seasonality is the main factor for the synchronization of different physiological classes (larvae, nymphs, and adults) in the population. Indeed, we notice that in cold climates, such as in the case of Canada, nymphs and adults stages are separated. Conversely, in Senegal where seasonal variations are almost nonexistent, due to low annual temperature fluctuation, activity periods for both tick stages are more clearly coexisting within the year. Tunisia, due to its Mediterranean climate, represents an intermediate context, where, according to our modeling approach, the three *H. scupense* stages whilst having separated peaks, continue to show a clear overlapping, which however appears more pronounced than recorded under field conditions [1, 8].

Furthermore, according to our model, at constant temperature, larvae are predicted to be present along the year at different densities and adults are expected to be present over a long period exceeding their usual activity season during summer.

We notice that, whatever the climate, all the physiological parameters of the life cycle of the ticks in our model are the same, except the lower bound of the temperature tolerance. This adaptation to the climate through a lower bound of the temperature tolerance could explain why some tick species are present in extreme climates (*e.g. Haemaphysalis longicornis,I.ricinus*) [39, 13]. We show that this adaptation can be done only on low values of developmental rate.

When applied to assess population dynamics, our model revealed, globally over the period 1995 − 2015 comparatively to the beginning of the century 1901 − 1925, a clear trend for increased tick densities in Canada with *I. cookie* and in Tunisia with *H. scupense*. Accordingly, temperature overall changes from 1901 − 25 to 1995 − 2015 are potentially more favorable to tick development in Canada and Tunisia, suggesting then *I. cookie* and *H. scupense* populations, may continue to expand if this dynamic of climatic change is maintained and if other tick populations regulating factors are not coming into play. Contrasting results were obtained in Senegal with the tick species *A. variegatum*. In fact, our simulations are showing that the *A. variegatum* population had known a clear two peaks during the last century, with a significant decrease in population densities between 1995 − 2005.

One of the advantages of our model is its incremental capacity to be extended to incorporate more accurate data on tick biology, and in particular survival of free stages to biotic and abiotic factors, if a more precise prospective analysis is required for a given tick species under specific environmental contexts. Furthermore, this advantage could allow its application to model pathogen transmission dynamics by predicting risk factors for vector-borne pathogens transmission and in particular vector abundance and seasonality, and by expressing the pathogen basic reproductive rate *R*_0_ according to (or as a function?) to *R*_0_ of the vector tick population.

In the current model, we only consider one climatic factor; temperature. Various biotic and abiotic elements interact continuously (e.g., relative humidity, hazardous chemicals, photoperiod, parasitism, etc.). As a result, populations can exhibit shifting responses to the environment, and some life stages will be more sensitive to these factors than others. For instance, a further study that examines a more explicit relationship between temperature and bioenergetic budgets dynamics, specifically in moulting ticks, will be required to establish a thorough understanding of the consequences of climate change on ticks’ life cycle.

## Acknowledgment

We thank Prof. Mohamed Gharbi for his valuable comments and assistance.

